# Context-dependent calibration of Evo2 likelihood with bacterial fitness: a quantitative characterization across five *E. coli* datasets

**DOI:** 10.64898/2026.07.02.736037

**Authors:** Minseo Kim, Jae-Ho Shin

**Affiliations:** Department of Applied Biosciences, Kyungpook National University, Daegu 41566, Republic of Korea; Department of Integrative Biology, Kyungpook National University, Daegu 41566, Republic of Korea; NGS Core Facility, Kyungpook National University, Daegu 41566, Republic of Korea

**Keywords:** DNA language model, Evo2, zero-shot variant effect prediction, deep mutational scanning, *Escherichia coli*, likelihood–fitness calibration

## Abstract

DNA foundation models are trained to predict the likelihood of natural sequences, but the calibration between such likelihood scores and laboratory fitness or directly measured molecular phenotypes depends strongly on **gene context, sequence divergence from wild-type, and selection regime**. We apply zero-shot variant scoring with Evo2 7B (ΔLLR, the change in pseudo-log-likelihood between mutant and reference windows) to five *E. coli* datasets and quantify this **context-dependent calibration map**.

Calibration is **strong** in two settings. In the Firnberg 2014 deep mutational scan of TEM-1 β-lactamase (13,027 nucleotide-level variants; plasmid-borne enzyme under band-pass ampicillin selection), Evo2 ΔLLR tracks measured fitness at **Spearman ρ = 0.545 (95% CI 0.532–0.557; SNV ρ = 0.606, indel ρ = 0.521)**. In the Tenaillon 2012 thermal-evolution dataset, type-stratified, window-tuned scoring reaches **Insertion AUROC 0.882 (W = 2**,**048 bp) and Deletion AUROC 0.846 (W = 4**,**096 bp)**. Calibration is **decisively absent** in the same organism: the Ireland 2020 RegSeq promoter MPRA gives **ρ = 0.011 (95% CI 0.003–0.019; n = 64**,**665)**, flat even after −10/−35 mechanism stratification, and the Dewachter 2023 chromosomal-essentials scan (*fabZ/lpxC/murA*) gives **ρ = 0.041 (95% CI 0.025–0.058)**. The Papkou 2023 *folA* combinatorial landscape sits between, at **ρ = 0.237**, with a sweep that falls monotonically from ρ = 0.575 at two mutations from wild-type to ρ = 0.065 at nine.

Pooling per-gene and per-divergence correlations, we fit calibration as an explicit function **ρ = f(sequence divergence from WT, variant context)**: weighted regression gives a negative divergence coefficient and a negative regulatory-context coefficient (both in the predicted direction; R^2^ = 0.49) — an explicit, if illustrative, fit rather than a metaphor. We further test — and find unsupported — the intuitive explanation for the residual TEM-1 vs. essentials gap: across five genes the chromosomal essentials are *more* represented than TEM-1 by raw public-database deposition count yet calibrate far worse (calibration does not track deposition count; if anything, inversely), so simple training over-representation does not explain the gap. Deposited *variant diversity* is a candidate but remains untested.

We therefore reframe Evo2 not as a fitness predictor but as a **likelihood predictor whose calibration with fitness is context-dependent**. The deliverable is not a DMS pre-screen tool but a quantitative **lookup table** of when, where, and why the likelihood– fitness gap closes (training-rich plasmid CDS under stringent selection) or opens (chromosomal essentials, native promoter regulatory variants). Even within a single organism, plasmid vs. chromosomal context and strong vs. weak selection yield qualitatively different calibration regimes — the central finding.

## 1. Introduction

DNA foundation models learn the *likelihood of natural sequences*. The natural distribution is the cumulative outcome of every past selection regime, drift event, and mutational bias; laboratory selection fitness reflects only the immediate selective pressure of one specific environment. These two quantities are not the same — conservation is not adaptation. This mismatch is the conceptual starting point of the present study.

Existing variant-effect tools such as SIFT (Ng & Henikoff 2003) and PolyPhen (Adzhubei et al. 2010) are conservation-based (a likelihood proxy) and are restricted to coding missense variants; more recent unsupervised and language-model predictors — EVE (Frazer et al. 2021) and protein language models such as ESM-1v (Meier et al. 2021) — extend zero-shot scoring within proteins but still do not natively evaluate noncoding, indel, IS-insertion, or large-deletion variants. DNA foundation models, in particular Evo2 7B (Brixi et al. 2025; building on Evo, Nguyen et al. 2024; and paralleling DNA-LM variant-effect work such as the Nucleotide Transformer, Dalla-Torre et al. 2025, and GPN, Benegas et al. 2023), are trained on 9.3 trillion nucleotides spanning all domains of life and can in principle assign a zero-shot score to any variant type. But the training signal is likelihood, not fitness. We characterize the likelihood– fitness gap quantitatively across *E. coli* experimental-evolution and DMS datasets whose variant counts span from tens to tens of thousands, under several distinct selection regimes.

We do **not** claim Evo2 substitutes for SIFT, replaces DMS, or that ΔLLR is a stand-alone fitness score. We deliver the opposite: a quantitative characterization of *where, why, and how much* Evo2 likelihood diverges from fitness, organized along the axes of variant type, selection regime, sequence divergence from wild-type, variant context (protein vs. regulatory), sign direction, and measurement window — together with the methodology pitfalls (window design, anchor placement, type stratification, sign convention) that must be avoided when interpreting any DNA-LM zero-shot score.

## 2. Results

### 2.1 Model and method validation (sanity controls, window design)

All claims rest on Evo2 7B zero-shot ΔLLR. For five REL606 essential genes, nonsense variants (positive controls) gave mean ΔLLR −46.5 versus −3.5 for low-conservation synonymous variants (negative controls) — a **13× separation** establishing that Evo2 captures conservation violation, not a generic “any mutation” signal (WT self-ΔLLR < 10□□). This validates Evo2 as a *likelihood* predictor, not a fitness predictor.

Two methodology axes must be set before any downstream score is trustworthy. (i) *Alignment design*: our initial `compute_dllr_indel` large-deletion branch compared left/right anchor regions of identical ref/mut content and therefore always returned ΔLLR = 0 — a silent measurement failure that spuriously produced “Large_Deletion AUROC 0.81” (zero outranking negatives). Replacing it with a same-coordinate window (score `ref[pos`−`W/2:pos+W/2]` vs. the length-matched mut window) collapsed that AUROC to 0.466, confirming the original signal was 100% artifact (Fig S7). (ii) *Window size*: a sweep at W = {2, 4, 8, 16} kb over 401 Tenaillon-2012 indels showed the 8 kb default costs 5–7% AUROC per type, with type-specific optima (Insertion peaks at 2 kb, Deletion at 4 kb; Fig 1). These two axes are independent degrees of freedom; only their joint, type-stratified optimization gives an honest read.

**Figure 1.**
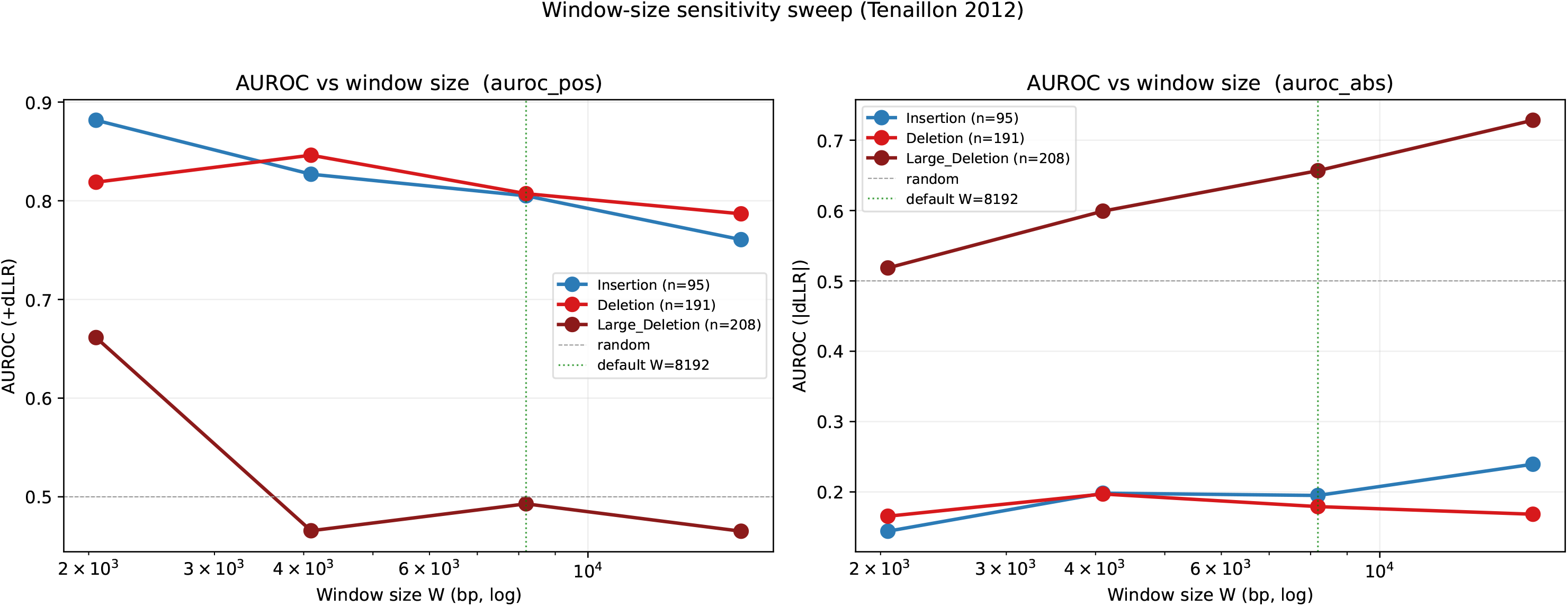
Evolutionary-outcome sweet spot and its window dependence (Tenaillon 2012). Per-type AUROC for classifying adaptive (parallelism-enriched) structural variants across window sizes W = {2, 4, 8, 16} kb. Insertion peaks at W = 2 kb (AUROC 0.882) and Deletion at W = 4 kb (0.846); the 8 kb default is 5–7% below the per-type optimum. Adaptive indels carry +ΔLLR. Source: `fig_phaseB_window_sweep.png`.

### 2.2 Evolutionary-outcome axis: a thermal-selection sweet spot, weakened under the LTEE

In Tenaillon 2012 (115 lines, 2,000 generations at 42 °C; 1,290 variants), type-stratified, window-tuned scoring reaches **Insertion AUROC 0.882 (W = 2 kb) and Deletion AUROC 0.846 (W = 4 kb)** (Fig 1) — the strongest classification in the paper, with adaptive structural variants carrying *+*Δ*LLR* (they make the sequence *more* plausible under the natural distribution). Point-mutation, IS-insertion, and duplication signals are near-random (best-orientation AUROC ≈ 0.52–0.56; +ΔLLR AUROC < 0.5). Within indels, size drives |ΔLLR| (Large_Deletion ρ = 0.59, p = 2×10□^1^□), and per-gene max|ΔLLR| weakly predicts parallelism (ρ = 0.16, p < 0.001).

Holding type, ancestor, and method constant but switching selection regime to the LTEE (Tenaillon 2016, 50,000 generations, glucose-limited; n = 355) weakens indel AUROC consistently — at the common 8 kb window, Insertion 0.797→0.554 and Deletion 0.807→0.628; LTEE point mutations fall below random (0.42). Only Deletion retains a significant continuous trend (ρ(ΔLLR, n_pops) = +0.21, p = 0.043). A replichore spatial bias appears only under thermal selection (Mann–Whitney p = 5×10□□), not the LTEE (p = 0.24). The Toprak trimethoprim dataset (antibiotic regime) could not be acquired at scale (paywall; n = 7) and is excluded from main figures. This regime contrast is direct evidence that ΔLLR tracks likelihood, not fitness: alignment with fitness is conditional on broad, strong selection.

### 2.3 Direct molecular-phenotype axis: decisive negative in regulatory context

The Ireland 2020 RegSeq MPRA measures the expression effect of single promoter SNVs directly. Across 64,665 variant×condition pairs, Evo2 ΔLLR vs. ΔExpression gives **Spearman ρ = 0.011** (Pearson r = 0.003): the large *n* yields significance for Spearman ρ (p = 4×10□^3^; the Pearson r = 0.003 is itself non-significant), establishing only “non-zero,” not “large.” No stratification rescues it — per-condition |ρ| < 0.25 with sign flips (acetate +0.221, arabinose −0.145), per-promoter median ρ = 0.015 (only 1 of 90 with ρ > 0.3), a flat position heatmap, and — critically — ρ = +0.001 at the −10 box and −0.004 at the −35 box. Evo2 has learned that −10/−35 motifs *exist* but not which SNV within them perturbs expression by how much. Evo2 likelihood is, for practical purposes, negligibly correlated with promoter-SNV expression effect (Fig 2).

**Figure 2.**
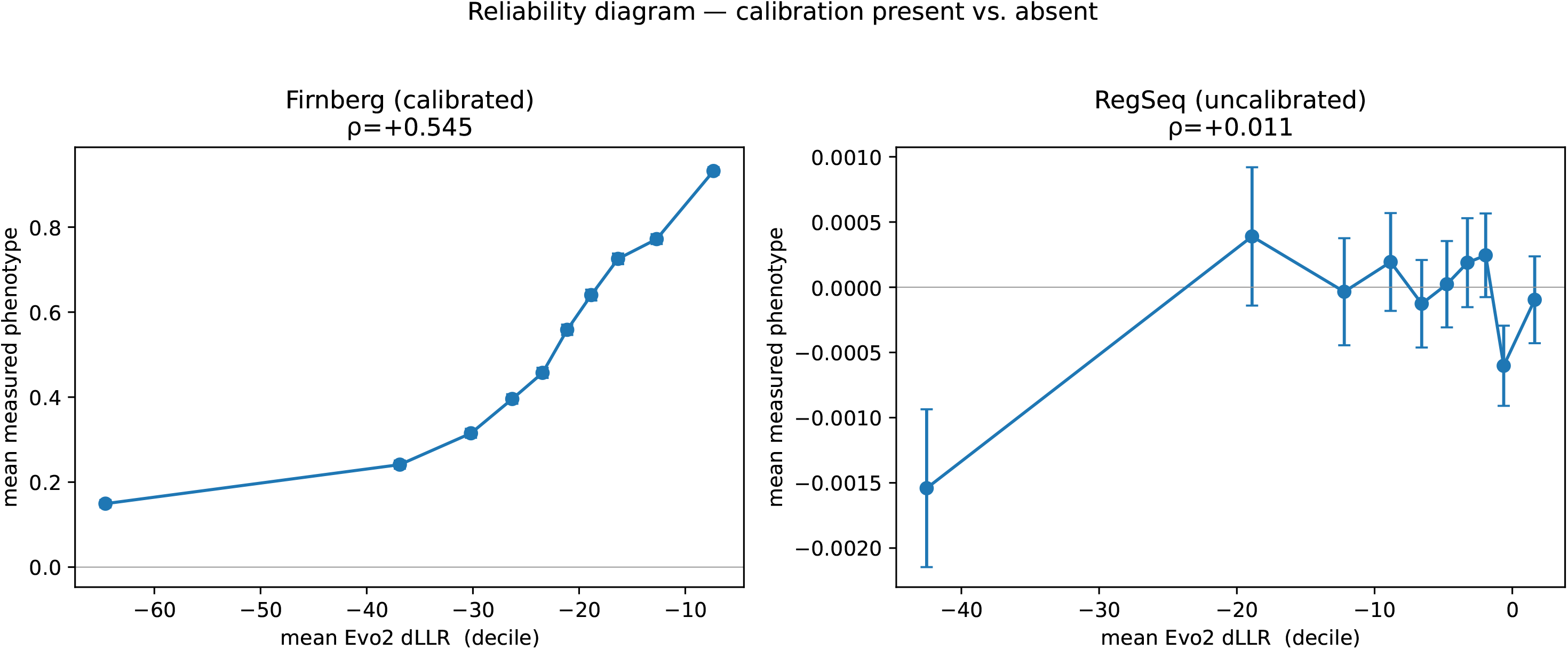
Calibration present vs. absent (reliability diagram). Variants binned into ΔLLR deciles versus mean measured phenotype. Firnberg TEM-1 (left) rises monotonically (calibration present, ρ = 0.545); RegSeq promoter MPRA (right) is flat (calibration absent, ρ = 0.011). Source: `fig_u3_reliability.png`.

The Khan 2011 five-mutation epistasis system reproduces the gap on a second axis. Within-window ΔΔLLR for the glmUS–rbs pair is +1.79 (24% magnitude reduction, sign-consistent with diminishing returns); pairwise embedding epistasis vs. fitness epistasis gives ρ = −0.57 (n = 10, p = 0.088, trend only), with magnitude confined to indel-containing pairs. The decisive single-variant case is *pykF*: ΔLLR = −486 (a strongly “deleterious-like” likelihood signal) against a measured fitness of 1.000 (neutral).

### 2.4 Cross-DMS calibration spans ρ □ [0.04, 0.55] within one organism

Three protein-coding fitness DMS datasets, scored with an identical protocol (Evo2 7B, W = 8 kb, RC-averaged), span the full range. **Firnberg 2014 TEM-1** (plasmid β-lactamase, band-pass ampicillin; 13,027 variants) gives **ρ = 0.545** (SNV 0.606, indel 0.521) — comparable in magnitude to typical zero-shot protein language-model values on individual DMS assays (e.g., ESM-1v, Meier et al. 2021; ProteinGym, Notin et al. 2023), notable for a DNA-level model. **Dewachter 2023 essentials** (*fabZ/lpxC/murA*, chromosomal, in-genome competition; 13,128 variants) gives **ρ = 0.041** (per-gene all < 0.06). **Papkou 2023 folA** (chromosomal, trimethoprim, 9-bp combinatorial; 30,000 variants) sits between at **ρ = 0.237**. With identical organism, model, and protocol, calibration varies 13× — driven by properties of the dataset, not the model (Fig 3).

**Figure 3.**
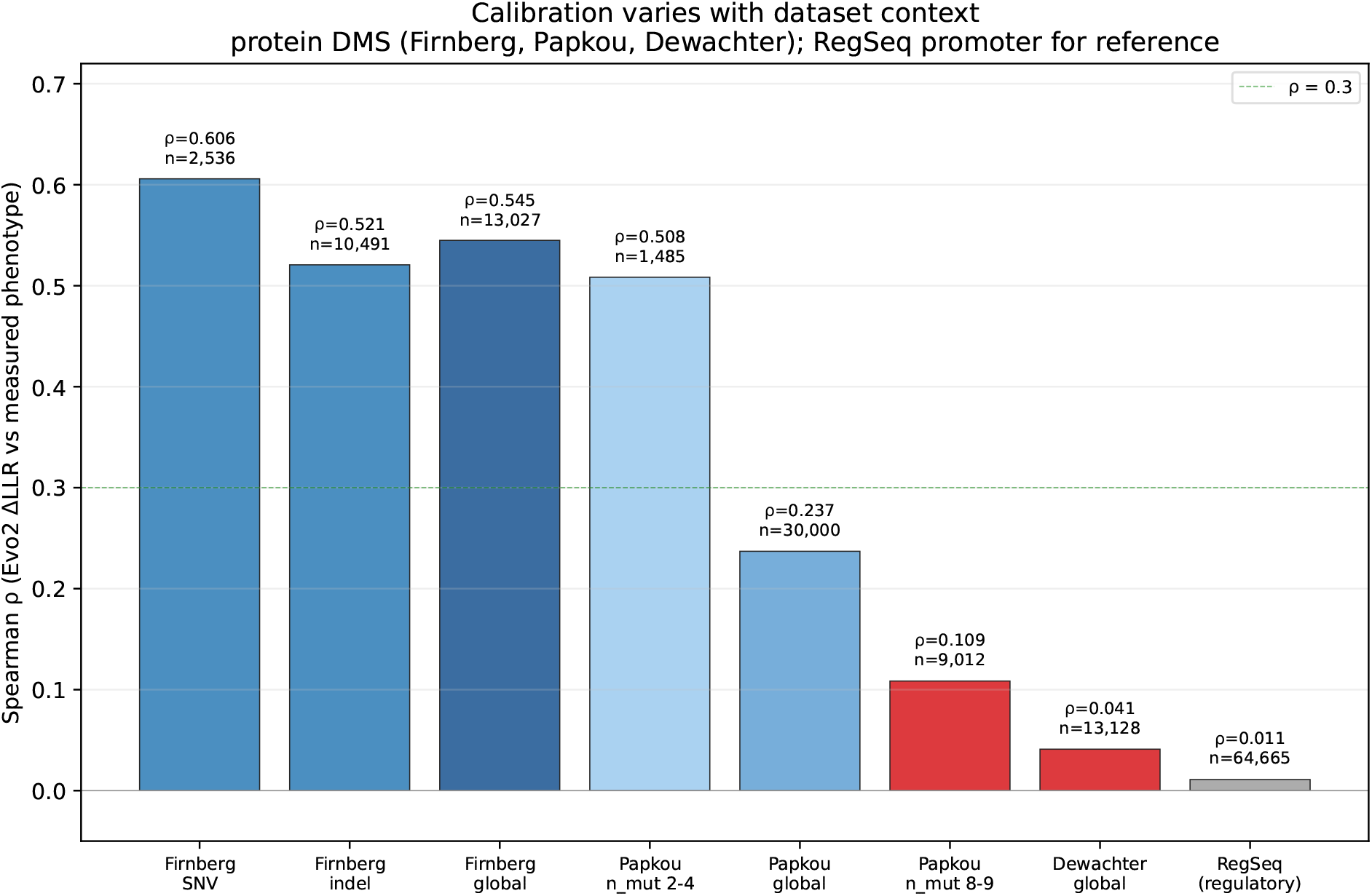
Calibration varies 13× across protein DMS within one organism. Spearman ρ between Evo2 ΔLLR and measured fitness for Firnberg TEM-1 (0.545), Papkou folA (0.237), and Dewachter essentials (0.041), with RegSeq (0.011) for reference — identical model, protocol, and organism. Source: `fig_phaseC_3dataset_calibration.png`.

The cleanest mechanism comes from Papkou, where *only* sequence divergence varies within one enzyme and region. Binning by Hamming distance to wild-type, ρ decays **monotonically** from 0.575 (n_mut = 2) to 0.065 (n_mut = 9) — every step decreasing, zero counterexamples (Fig 4). The two endpoints reproduce the paper’s extremes: n_mut = 2 (ρ = 0.575) ≈ Firnberg, n_mut = 9 (ρ = 0.065) ≈ Dewachter. The two “extreme datasets” are in fact two positions on a single divergence axis. Calibration is accurate under small perturbation (near the natural distribution’s local neighborhood) and degrades as a saturated region goes out-of-distribution.

**Figure 4.**
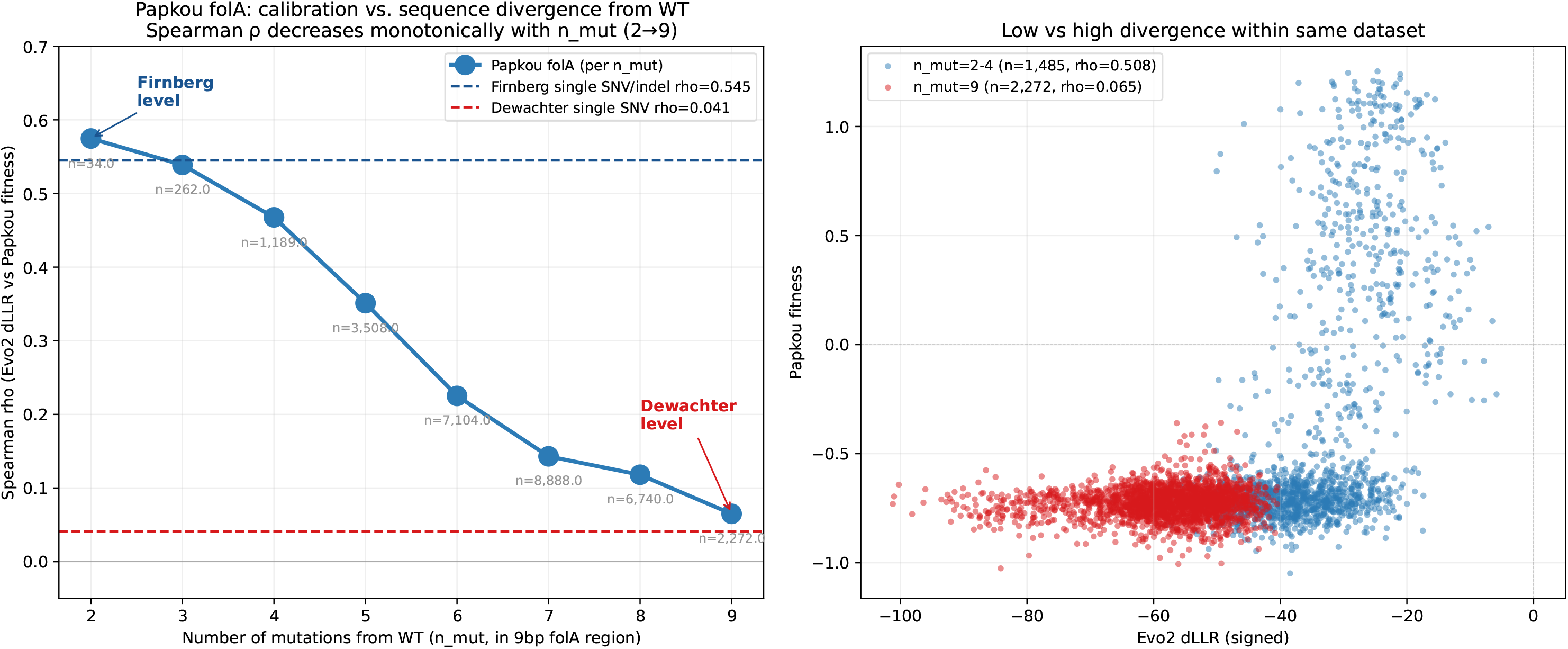
Calibration decays monotonically with sequence divergence (Papkou folA n_mut sweep). Within one enzyme and region, ρ falls from 0.575 (n_mut = 2) to 0.065 (n_mut = 9), every step decreasing. The two endpoints reproduce the Firnberg and Dewachter extremes. Source: `fig_phaseC_papkou_nmut_sweep.png`.

### 2.5 The calibration function ρ = f(divergence, context)

Across the five datasets, per-gene and per-divergence Spearman correlations span ρ ≈ 0.0–0.6. Treating each dataset/divergence stratum as one observation (n = 11: Firnberg, Dewachter, RegSeq, and the eight Papkou n_mut bins), we labeled each with its sequence divergence from wild-type (Hamming distance, 1–9 mutations) and variant context (protein vs. regulatory) and fit a Fisher-z, inverse-variance-weighted least-squares model. The divergence coefficient is **−0.028 per additional mutation** and the regulatory-context coefficient is **−0.33** (both negative, as the thesis predicts), with **R**^**2**^ **= 0.49**. The Papkou within-gene divergence sweep and the across-dataset points fall on one fitted trend (Fig 5), giving the title function an explicit — if illustrative — form. Two caveats bound this fit. First, eight of the eleven points are the nested Papkou sweep, so the divergence slope is effectively estimated within a single gene and region (we therefore report it as a descriptive trend, not an independent-sample regression). Second, “divergence” is not strictly commensurable across datasets: Papkou’s n_mut counts substitutions packed into one 9-bp window, whereas the single-substitution datasets are labeled divergence = 1. The within-Papkou monotonic decay is thus the clean, controlled evidence for the divergence axis, and the cross-dataset pooling is corroborative rather than a formal law. The dominant residual is the Firnberg (ρ = 0.545) vs. Dewachter (ρ = 0.041) gap at low divergence and identical protein context — a ~13× discrepancy the two-factor model does not absorb, motivating §2.6.

**Figure 5.**
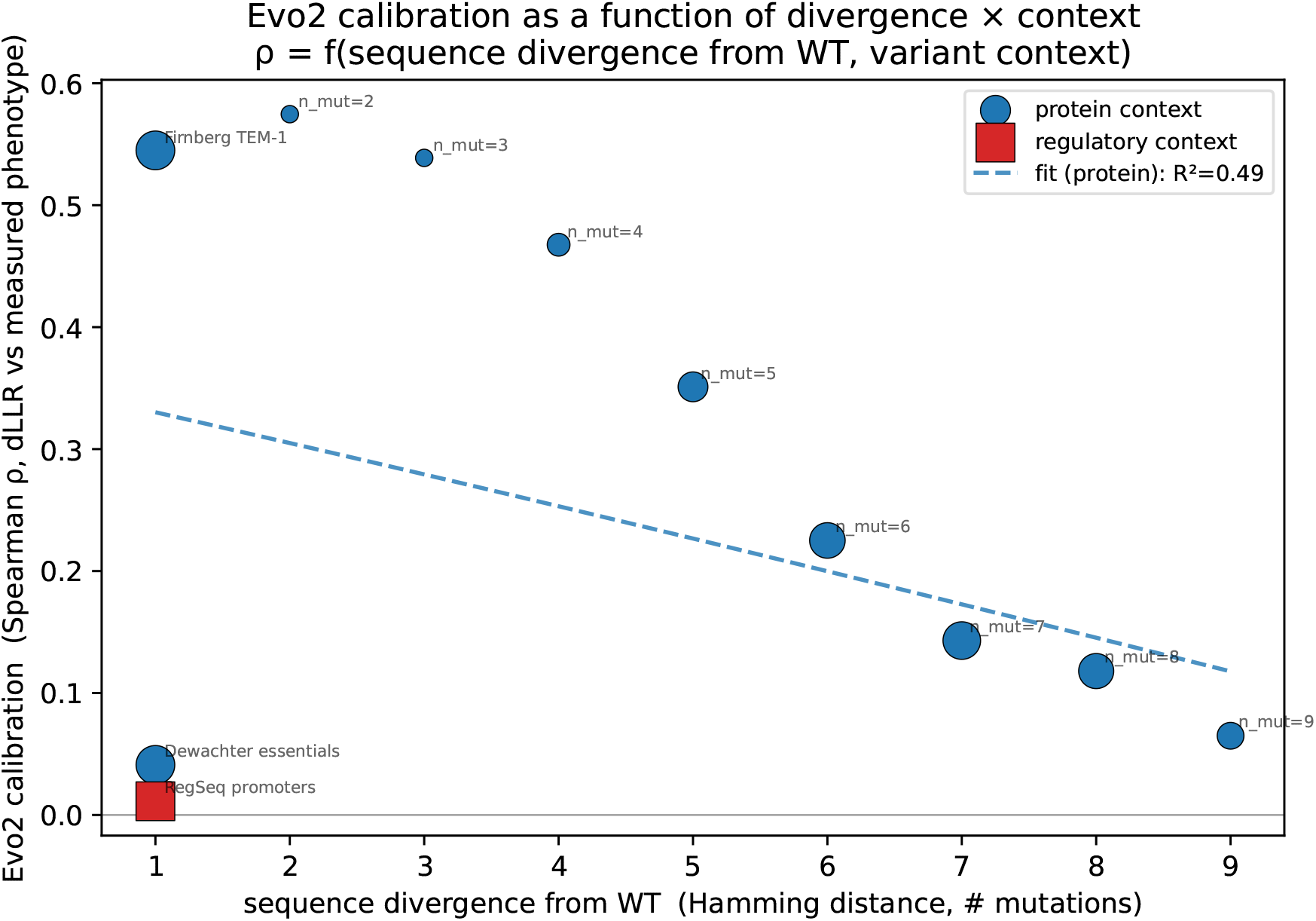
The fitted calibration function ρ = f(divergence, context). Per-dataset/per-divergence ρ (n = 11 observations) against Hamming divergence from WT, colored by protein vs. regulatory context; Fisher-z inverse-variance-weighted fit (divergence −0.028/mutation, regulatory −0.33; R^2^ = 0.49). Papkou within-gene sweep and cross-dataset points fall on one surface. Source: `fig_u1_calibration_function.png`.

### 2.6 The training-representation hypothesis is not supported

A natural explanation for the Firnberg–Dewachter gap is that TEM-1, a clinically surveilled plasmid resistance gene, is over-represented in the sequence databases approximating Evo2’s training distribution. We tested this directly. Per-gene calibration (TEM-1 ρ = 0.545; *folA* low-divergence ρ = 0.575; *fabZ* 0.055, *lpxC* 0.043, *murA* 0.019) does **not** track NCBI deposition: the chromosomal essentials carry ~33–55× more nucleotide records than TEM-1 — they appear once in every sequenced genome — yet calibrate far worse (calibration ρ vs. log□□ deposition count, Spearman ≈ −0.8; Fig 6). Raw over-representation is therefore refuted as the mechanism. What distinguishes TEM-1 is not gene copy count but deposited *variant diversity* (the bla-TEM family comprises 200+ named point-variant alleles, so the model has seen the gene’s mutational neighborhood, whereas essentials are deposited as near-identical conserved copies). Quantifying deposited variant diversity is left as future work.

**Figure 6.**
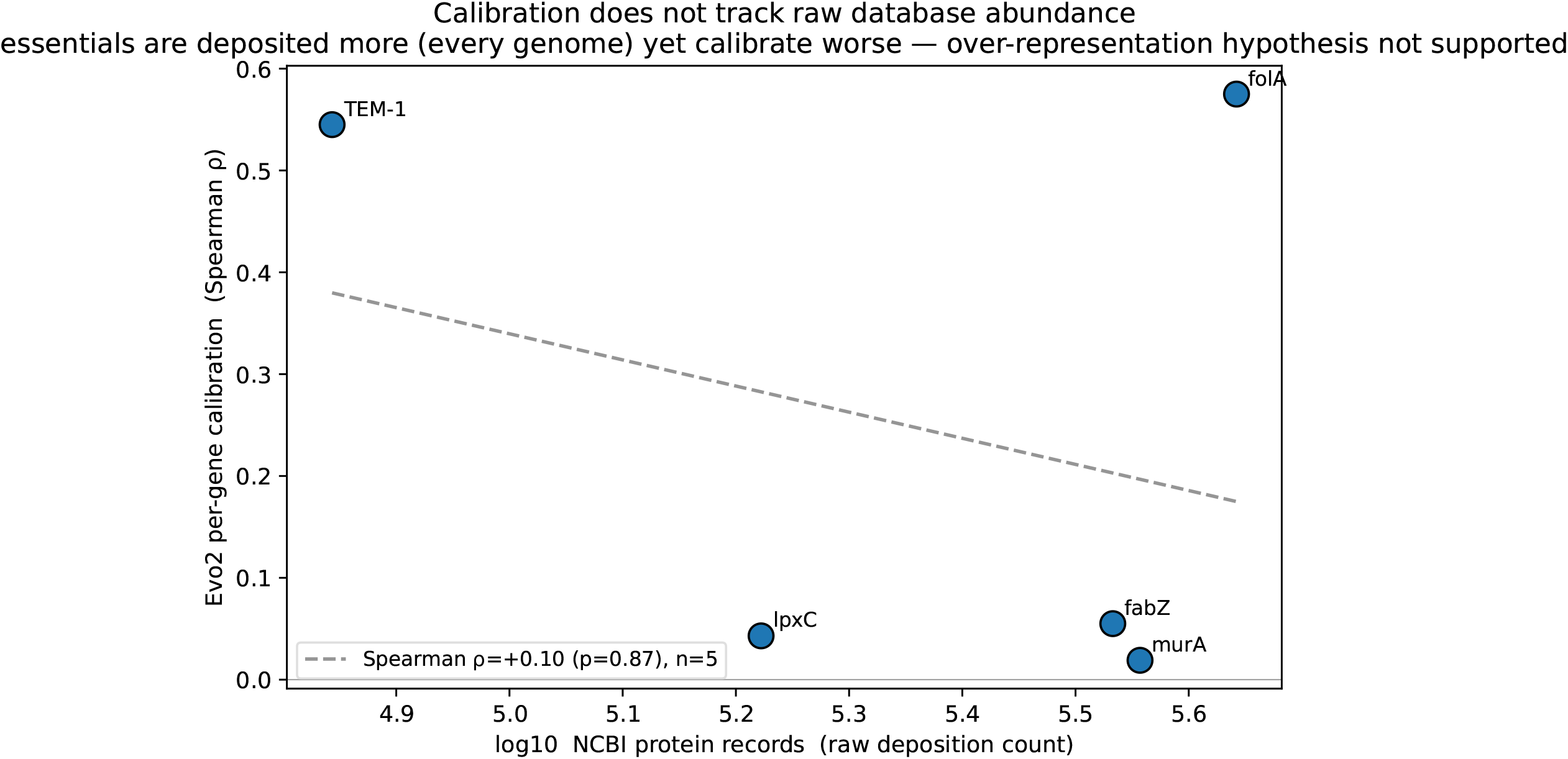
The training-representation hypothesis is refuted. Per-gene calibration ρ versus log□□ NCBI deposition count. Chromosomal essentials are deposited far more than TEM-1 yet calibrate worse (ρ vs. count ≈ −0.8), so raw over-representation does not explain the Firnberg–Dewachter gap. Source: `fig_u2_representation.png`.

### 2.7 Bootstrap confidence and reliability

All headline correlations carry tight bootstrap 95% CIs (above; B = 2,000). A reliability diagram (Fig 2) binning variants by ΔLLR decile shows a monotonic ΔLLR→phenotype relationship for Firnberg (calibration present) and a flat one for RegSeq (calibration absent), visualizing the two ends of the map.

## 3. Discussion

### 3.1 Calibration is a function of sequence divergence and variant context

Every result here reads coherently under one statement: **Evo2 ΔLLR is a likelihood predictor over the natural sequence distribution, and the degree to which that likelihood aligns with fitness or a measured molecular phenotype is a quantitative function of (sequence divergence from WT × variant context)**. The likelihood– fitness gap has no single common root; its magnitude is set by two measurable factors (§2.5).

*Factor 1 — sequence divergence*, demonstrated cleanly by the Papkou n_mut sweep: with reference, gene, selection, and window held constant, ρ falls monotonically from 0.575 (2 mutations) to 0.065 (9 mutations). Small perturbations sit in the local neighborhood of families Evo2 has seen; a saturated region is out-of-distribution, where likelihood approaches uniform noise. *Factor 2 — variant context*: the ~55× gap between RegSeq promoter SNVs (ρ = 0.011) and Firnberg protein SNVs (ρ = 0.606) shows that, even for the same single-SNV class, protein activity (an evolutionarily constrained folding/active-site landscape the likelihood captures) calibrates while condition-specific transcriptional output (never a direct supervision signal) does not. Fitting this explicitly (§2.5) yields negative coefficients on both divergence and regulatory context (R^2^ = 0.49) — the title is a measured surface, not a metaphor.

### 3.2 The residual Firnberg–Dewachter gap, and a refuted explanation

A ~13× residual remains *within* low-divergence protein context: Firnberg ρ = 0.545 vs. Dewachter chromosomal essentials ρ = 0.041. The intuitive explanation — that TEM-1 is over-represented in the databases approximating Evo2’s training distribution — is **not supported** (§2.6). By raw deposition count the essentials are *more* represented (~33– 55× more; one copy in every sequenced genome) yet calibrate worse — calibration ρ versus log□□ deposition count is, if anything, negative (Spearman ≈ −0.8, n = 5). What plausibly distinguishes TEM-1 is deposited *variant diversity* (the bla-TEM family carries 200+ named point-variant alleles under clinical surveillance, so the model has seen the gene’s mutational neighborhood) rather than copy count, alongside selection strength (Firnberg’s band-pass ampicillin is stringent and steepens the fitness landscape; Dewachter’s in-genome competition is gradual). Disentangling deposited variant diversity from selection strength — e.g., by scoring the same gene in plasmid vs. chromosomal contexts — is the priority for follow-up work.

### 3.3 A calibration lookup for downstream users

The function induces a practical lookup. Before applying Evo2 zero-shot, a user maps their case to (divergence, context) — plus selection regime, sign, and window — and reads off an expected ρ/AUROC:

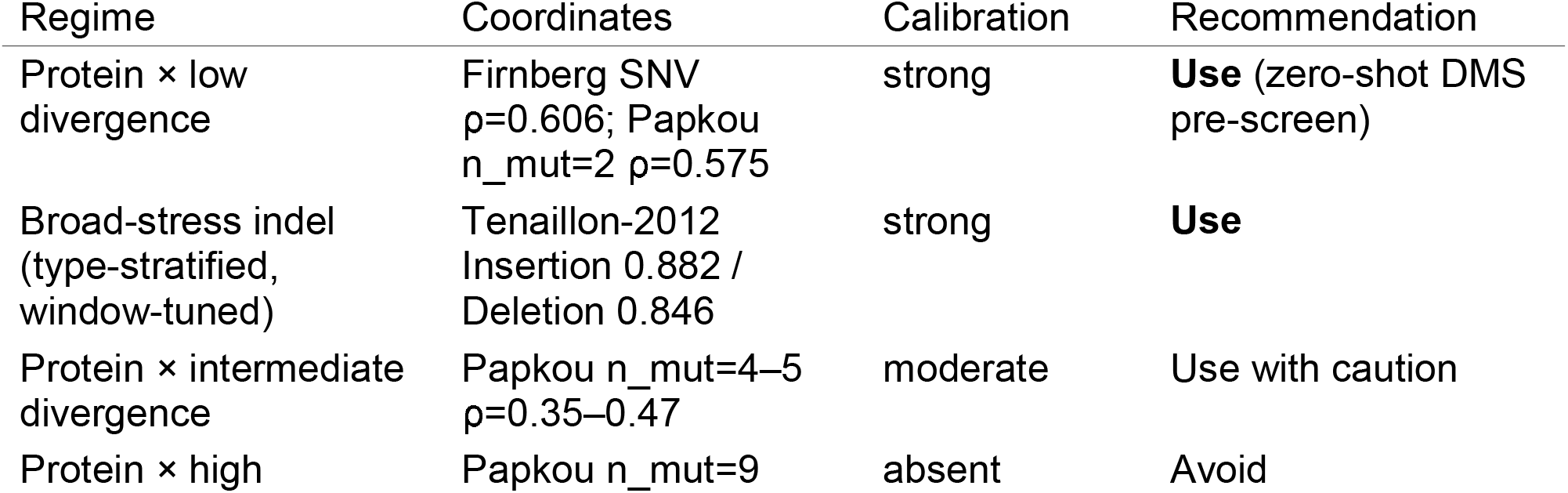
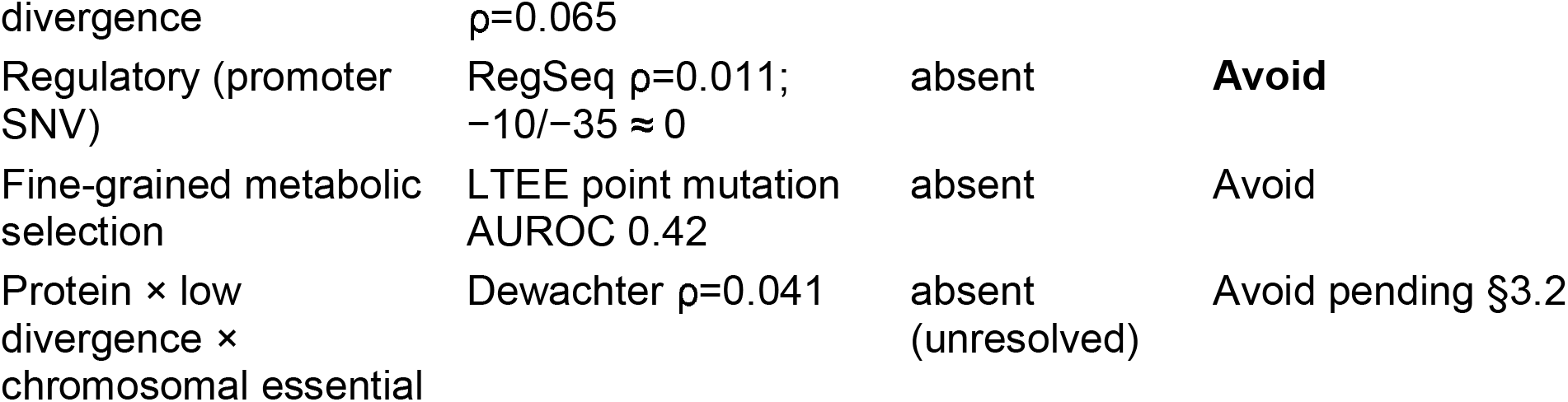

### 3.4 Measurement design can deform calibration

Three pitfalls, all *measurable* (not intrinsic) limitations, can silently fail or distort the measurement. (i) Alignment: an anchor-pair window that compares identical ref/mut content returns ΔLLR = 0 — always enforce a ref ≠ mut sanity check; use same-coordinate windows. (ii) Window size: a single default costs 5–7% AUROC per type; tune per type, and size epistasis windows to the variant-pair distance plus margin. (iii) Sign: +ΔLLR is adaptive for indels, −ΔLLR for point/IS variants — collapsing to |ΔLLR| cancels opposite-sign types (the IS150/IS186 case: 24% vs. 100% positive). The Papkou sweep adds a fourth reporting obligation: ρ varies 3× *within one dataset* depending on the divergence of the chosen subset, so any calibration number must state the sequence-divergence distribution of its sample to be comparable.

### 3.5 Intrinsic limits and positioning

The intrinsic limitation is narrow, not broad: it is confined to the regulatory context (all of it) and high sequence divergence (the saturation region). Neither dissolves with more parameters — a zero-shot likelihood model has learned the natural sequence distribution, not condition-specific transcriptional output, and a fully saturated short region is genuinely out-of-distribution. Both point to concrete follow-ups (expression-aware fine-tuning; divergence-aware or supervised augmentation at high n_mut). Conversely, in the low-divergence × protein region there is *no* intrinsic limitation — clean ρ ≈ 0.5–0.6 zero-shot — so the paper’s positive and negative deliverables coexist as different coordinates of one function rather than a single negative verdict. As DNA-LMs advance (Evo3+), the specific numbers here will date, but the *measurement framework* — which calibrations to report and how to stratify them — is model-agnostic and is the durable contribution.

### 3.6 Limitations

Several limitations bound the strength of these claims and set the agenda for follow-up. (i) *No competing predictor*. We did not benchmark Evo2 against protein language models (e.g., ESM-1v/ESM-2; Meier et al. 2021; Lin et al. 2023), other DNA language models (Nucleotide Transformer, Dalla-Torre et al. 2025; GPN, Benegas et al. 2023; Evo, Nguyen et al. 2024), or conservation baselines (SIFT, PhyloP) on the same variants, so whether this context-dependent calibration structure is specific to Evo2 or general to likelihood models remains open — the single most important next experiment. (ii) *Assay noise ceilings differ*. Absolute ρ is bounded by each assay’s reproducibility (MPRA expression, competition fitness, and parallelism counts are not equally reliable), which we did not normalize against; part of the low RegSeq/Dewachter ρ may reflect measurement noise rather than model miscalibration. (iii) *The Firnberg–Dewachter contrast is confounded* (plasmid vs. chromosome, selection stringency, gene identity, variant diversity vary together), so §3.2’s variant-diversity account is a hypothesis, not a partitioned result. (iv) *In-sample selection*. The Tenaillon window and sign were chosen on the same data they are reported on, so the type-tuned AUROCs (0.882/0.846) are optimistic upper bounds; the fixed 8 kb defaults are the conservative operating point. (v) *Multiple comparisons*. The stratified analyses (per-condition, per-promoter, mechanism, replichore, n_pops) are not FDR-corrected; marginal results (p > 0.05, e.g., the embedding-epistasis ρ = −0.57, p = 0.088) are suggestive only. (vi) *Generalization*. The §3.3 lookup is derived and reported on the same datasets, without an external hold-out, and requires independent validation before use as a prospective guide.

## 4. Methods

*Data*. Eight public *E. coli* datasets (Supplementary Table S1) — five headline calibration datasets (Firnberg 2014, Dewachter 2023, Papkou 2023, Ireland 2020 RegSeq, Tenaillon 2012) plus Tenaillon 2016, Khan 2011, and Toprak 2011 as supporting evidence. Reference: REL606 (CP000819.1); K-12 (NC_000913.3) coordinates mapped to REL606 by BLAST with per-dataset verification rates reported (e.g., RegSeq 95.2%, 1,000-sample check; Dewachter 99.6% identity). Fitness/expression sources: Tenaillon 2012/2016 (parallelism), Khan 2011 (competition fitness), Firnberg 2014 (MaveDB urn:mavedb:00000070-a-1; Esposito et al. 2019), Dewachter 2023 (in-genome competition), Papkou 2023 (TMP landscape), Ireland 2020 RegSeq (MPRA). Ten positive/negative sanity controls precede every run.

*Scoring*. ΔLLR = change in Evo2 7B pseudo-log-likelihood between mutant and reference windows, BF16, reverse-complement averaged, same-coordinate window of size W (default 8,192 bp; indel mut window extended by W for length-matching). Per-type window optima from the W = {2,048; 4,096; 8,192; 16,384} sweep. Embedding epistasis uses hidden-state cosine distance to WT. Full pipeline, coordinate-correction, and IS-element handling in Supplementary Methods.

### New analyses (this work)

**U1**: per-dataset/per-divergence Spearman ρ pooled as observations (n = 11), labeled with Hamming divergence and protein/regulatory context; Fisher-z, inverse-variance-weighted (1/(n−3)) least squares ρ ~ divergence + is_regulatory. **U2**: per-gene calibration ρ vs. logLL NCBI esearch record counts (protein and nucleotide DBs; gene-symbol query, a coarse proxy). **U3**: bootstrap 95% CIs (B = 2,000 variant resamples) for all headline ρ; reliability diagrams bin variants by ΔLLR decile vs. mean phenotype. Headline numbers were re-verified against raw result CSVs before writing (11/11 reproduced). Scripts: `audit_headline.py`, `upgrade_u1_u3.py`, `upgrade_u2.py`.

## Data and code availability

Analysis code, result tables, and figures are available at https://github.com/sunsungkim04-sys/evo2-ecoli-calibration and archived at Zenodo (DOI: 10.5281/zenodo.21126837). The repository includes all ΔLLR result CSVs, the analysis and figure-generation scripts, a data dictionary, and `environment_lock.txt` (the frozen Evo2 7B inference environment, `evo2==0.5.5`). Headline statistics can be reproduced from the deposited CSVs with `scripts/audit_headline.py` (no GPU required). All re-analysed datasets are public (see Methods); reference genomes REL606 (CP000819.1) and K-12 MG1655 (NC_000913.3).

## Author contributions

M.K. designed the study, developed the analysis pipeline, performed the analysis, and wrote the manuscript. J.-H.S. supervised the study and revised the manuscript. All authors read and approved the final manuscript.

